# Dynamics of visual perceptual decision-making in freely behaving mice

**DOI:** 10.1101/2020.02.20.958652

**Authors:** Wen-Kai You, Shreesh P. Mysore

## Abstract

Studying the temporal dynamics of perceptual decisions offers key insights into the cognitive processes contributing to it. Conducting such investigation in a genetically tractable animal model can facilitate the subsequent unpacking of the mechanistic basis of different stages in perceptual dynamics. Here, we investigated the time course as well as fundamental psychophysical constants governing visual perceptual decision-making in freely behaving mice. We did so by analyzing response accuracy against reaction time (i.e., conditional accuracy), in a series of 2-AFC orientation discrimination tasks in which we varied target size, luminance, duration, and presence of a foil. Our results quantified two distinct stages in the time course of mouse visual decision-making - a ‘sensory encoding’ stage, in which conditional accuracy exhibits a classic tradeoff with response speed, and a subsequent ‘short term memory-dependent’ stage in which conditional accuracy exhibits a classic asymptotic decay following stimulus offset. We estimated the duration of visual sensory encoding as 200-320 ms across tasks, the lower bound of the duration of short-term memory as ~1700 ms, and the briefest duration of visual stimulus input that is informative as ≤50 ms. Separately, by varying stimulus onset delay, we demonstrated that the conditional accuracy function and RT distribution can be independently modulated, and found that the duration for which mice naturally withhold from responding is a quantitative metric of impulsivity. Taken together, our results establish a quantitative foundation for investigating the neural circuit bases of visual decision dynamics in mice.

**SIGNIFICANCE STATEMENT:** This study presents a quantitative breakdown of the time course of visual decision-making in mice during naturalistic behavior. It demonstrates parallel stages in mouse visual perceptual decision dynamics to those in humans, estimates their durations, and shows that mice are able to discriminate well under challenging visual conditions – with stimuli that are brief, low luminance, and small. These results set the stage for investigating the neural bases of visual perceptual decision dynamics and their dysfunction in mice.

## INTRODUCTION

Exploring the temporal dynamics of perceptual decisions from onset of the sensory input through the initiation of behavioral responses affords a key window into the underlying cognitive processes [1–3]. Investigations of such dynamics in humans [4, 5], and other species[6–8] have revealed distinct stages in perceptual processing, their timing, and their interactions. [9–11]. Performing such investigations in a genetically tractable animal model can additionally facilitate the subsequent unpacking of the mechanistic basis of different stages in perceptual dynamics. However, despite the recent rise in the use of the laboratory mouse for the study of the visual system [12–14] and of visually guided decision-making [15–25], the temporal dynamics of visual perceptual decisions represents a significant gap in mouse visual psychophysics [26–28].

In this study, we adapted approaches from human psychophysical studies to investigate the dynamics of visual decision-making in freely behaving mice. In a series of experiments involving touchscreen-based [24, 29], 2-alternative forced choice (2-AFC) orientation discrimination tasks, we investigated the effect of stimulus size, luminance, duration, delay, and the presence of a competing foil on mouse decision performance (accuracy and reaction time), and importantly, on the conditional accuracy function. We identified two distinct stages in the time-course of mouse visual decision-making within a trial, as has been reported in humans [30–37]. In the first ‘sensory encoding’ stage [30–33], response accuracy exhibited a classic tradeoff with response speed, and asymptoted to a peak level. In the next stage, response accuracy did not exhibit such a tradeoff, but instead, decayed following stimulus offset, consistent with a classic short-term memory (STM)-dependent process [34–37]. Combining these results with those from drift diffusion modeling [38] allowed us to estimate fundamental psychophysical constants in mouse perceptual decision-making: the time needed by mice to complete visual sensory encoding, the duration for which their short term memory can intrinsically support discrimination behavior after stimulus input is removed, and the shortest visual stimulus duration that is informative. Additionally, by varying stimulus onset delay, we demonstrated that the two components of accuracy, namely, the conditional accuracy function and the RT distribution can be independently modulated by task parameters. This also allowed a quantitative estimation of impulsivity of mice. Together, this study reveals parallels between mouse and human visual decision dynamics, despite differences in their sensory apparatuses, and enable investigations into the neural circuit underpinnings of the time course of perceptual decision-making in mice.

## METHODS

### Animals

Thirty-seven mice (33 C57B16/J mice, all male; 4 PV-Cre mice, 3 female, Jackson Labs) were housed in a temperature (~75F) and humidity (~55%) controlled facility on a 12:12h light:dark cycle; ZT0=7 am. All procedures followed the NIH guidelines and were approved by the [Author Institutions] Animal Care and Use Committee (ACUC). Animals were allowed to acclimate for at least one week, with *ad libitum* access to food and water before water regulation was initiated per previously published procedures [39]. Briefly, mice were individually housed (for monitoring and control of daily water intake of each identified animal), and administered 1mL water per day to taper their body weight down, over the course of 5-7 days, to 80-85% of each animal’s free-feeding baseline weight. During behavioral training/testing, the primary source of water for mice was as a reinforcer for correct performance: 10 μL of water was provided for every correct response. Experiments were all carried out in the light phase.

### Apparatus

Behavioral training and testing were performed in soundproof operant chambers equipped with a touchscreen (Med Associates Inc.), a custom-built reward port (fluid well), infrared video cameras, a house light and a magazine light above the reward port. The reward port was located at the opposite wall of the chamber relative to the touchscreen (Fig. 1A, 1-1A). Mice were placed within a clear plexiglass tube (5cm diameter) that connects the touchscreen and the reward port. A thin plexiglass mask (3 mm thickness) was placed 3 mm in front of the touchscreen with three apertures (1cm diameter) through which mouse was allowed to interact with the screen via nose-touch. The ‘left’ and ‘right’ apertures were placed 3cm apart (center-to-center) along the base of the triangle, and a ‘central’ aperture, at the apex of the triangle, was 1.5 cm below the midpoint of the base. All experimental procedures were executed using control software (K-limbic, Med-Associates).

**Figure 1.**
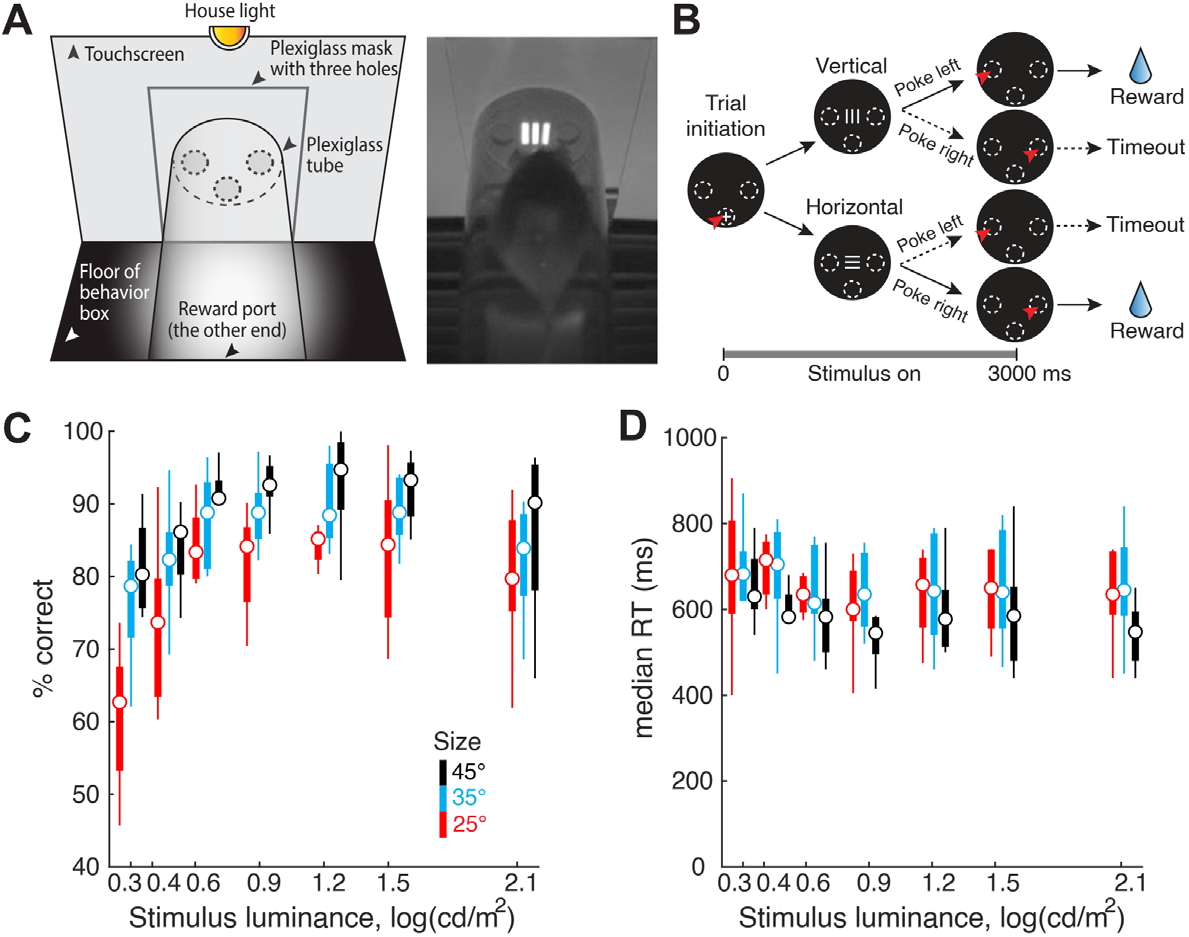
Stimulus contrast and size modulate orientation discrimination performance in freely behaving mice. **(A)**<u>Left</u>: Schematic of touchscreen-based experimental setup showing key components. <u>Right</u>: Snapshot of freely behaving mouse facing a visual stimulus on the touchscreen. **(B)** Schematic of 2-AFC task design. Black discs: Screenshots of touchscreen with visual stimuli; dashed ovals: locations of holes through which mice can interact with touchscreen; white ‘+’: zeroing cross presented within central response hole at start of each trial; red arrowhead: nose-touch by mouse. Shown also are vertical or horizontal grating stimuli, and reinforcement (water)/punishment (timeout) schedule. <u>Bottom</u>: Trial timeline. 0 ms corresponds to the instant at which the mouse touches the zeroing cross (trial initiation). Immediately following this, the target grating was presented and stayed on for 3s, or until the mouse responded, whichever came first. Vertical and horizontal targets were interleaved randomly. **(C)** Psychometric plots of discrimination accuracy against stimulus luminance (n=8 mice). Different colors correspond to different target sizes. 2-way ANOVA, p<0.001 (luminance), p<0.001 (size), p=0.498 (interaction). Effect size η^2^=0.292 (luminance), η^2^=0.192 (size), η^2^=0.037 (interaction). For each stimulus size/luminance, the box plot shows the median (the central mark), and the 25^th^ and 75th percentiles (the bottom and top edge of the box) of the group (n=8). The whiskers extend to the most extreme data points not considered as outliers. **(D)** Plot of median reaction time (RT) against stimulus contrast. 2-way ANOVA, p=0.998 (contrast), p=0.004 (size), p=1 (interaction). Effect size η^2^=0.003 (luminance), η^2^=0.071 (size), η^2^=0.010 (interaction). **See also Fig. 1-1.**

### Visual stimuli

Visual stimuli were bright objects on the dark background (luminance = 1.32 cd/m^2^). A small cross (60×60 pixels; luminance = 130 cd/m^2^) was presented in the central aperture and had to be touched to initiate each trial. Oriented gratings (horizontal or vertical) were generated using a square wave, with fixed spatial frequency (24 pixels/cycle) known to be effective for mice to discriminate [17]. The dark phase of the grating was black, identical to the background (luminance, L_dark_= 1.32 cd/m^2^), and the bright phase was varied between 1.73 cd/m^2^ and 130 cd/m^2^ depending on the tasks (see below). –The size of the stimulus was also varied depending on the task, ranging from 60 pixels × 60 pixels to 108 pixels × 108 pixels, which subtended 25-45 visual degrees at a viewing distance of 2 cm from the screen (Fig. 1-1A).

### Experimental procedure and behavioral training

Each mouse was run for one 30 min behavioral session per day, with each session yielding 80-180 trials. Each trial in a session was initiated by the mouse touching the zeroing cross. Upon trial initiation, the cross vanished, and the visual stimulus (or stimuli) were immediately presented (except in the delay task), for a duration of 0.1-3s depending on the task (see below). Mice were trained to report the orientation of target grating, by nose-touching the correct response aperture (vertical →; left; horizontal → right). A correct response triggered a tone (600 Hz, 1 sec), the magazine light turning on, and the delivery of 10μL of water. When mice turned to consumed the reward, their head entry into the reward port was detected by an infrared sensor which caused the zeroing cross (for the next trial) to be presented again. An incorrect response triggered a 5-s timeout, during which the house light and the magazine light were both on and zeroing cross was unavailable for the next trial to be initiated. A failure to respond within 3s (starting stimulus presentation) resulted in a trial reset: the stimulus vanished and the zeroing cross was presented immediately (without a timeout penalty), to allow initiation of the next trial. Well-trained animals failed to respond on fewer than 5% of the total number of trials, and there were no systematic differences in the proportion of such missed trials between different conditions. Within each daily 30-minute behavioral session, mice consumed approximately 1mL of water. If a mouse failed to collect enough water from the behavioral session, they were provided with a water supplement using a small plastic dish in their home cage.

### Single-stimulus discrimination task

Upon trial initiation, a single grating stimulus (i.e., the ‘target’) was presented above the central aperture, at the same horizontal level as the left and right apertures, and mice were required to report its orientation with the appropriate nose-touch (Fig. 1B). When stimulus size and luminance were manipulated (Fig. 1, and 2), three different sizes were tested: 60×60, 84×84, 108×108 (pixels × pixels). For each size, seven different levels of luminance were tested: 2.00, 2.59, 4.37, 7.55, 16.2, 34.3, 130 cd/m^2^. (These corresponded nominally to Michelson’s contrasts of 20%, 32%, 54%, 70%, 85%, 93%, 98%, respectively; Michelson’s contrast is computed as (luminance_bright_ - luminance_dark_) / (luminance_bright_ + luminance_dark_) *100.) Trials with different stimulus luminance at a particular size were interleaved randomly throughout a session, while trials with different stimulus sizes were examined on different days. When the stimulus duration was manipulated (Fig. 3), the luminance (130 cd/m^2^) and size (60 pix × 60 pix) of the grating were fixed, and eleven different stimulus durations were tested: 100 ms, 200, 300, 400, 500, 600, 800, 1000, 1500, 2000, 3000 ms. The stimulus duration was fixed for a given day, and across days, was varied in a descending sequence from 3000 ms to 100 ms. When the stimulus onset delay was manipulated (Fig. 5), the luminance (130 cd/m^2^), size (60 pix × 60 pix), and duration (600 ms) of the grating were fixed. Three different delays were tested: 0, 100, and 200 ms. The delay duration was fixed for a given day, and varied in an ascending sequence from 0 ms to 200 ms.

**Figure 2.**
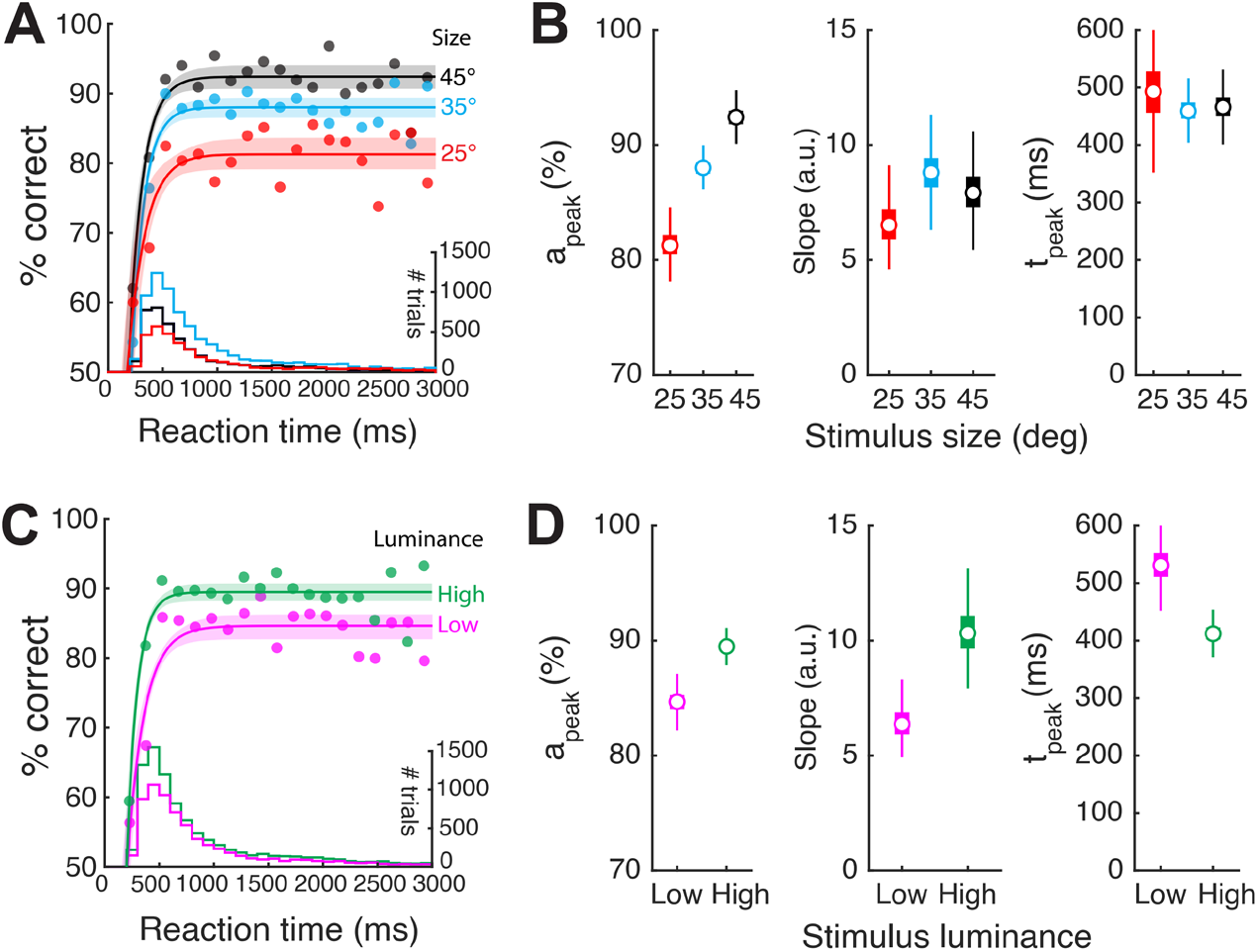
Stimulus size and luminance modulate the sensory encoding stage of the conditional accuracy function (CAF). **(A)** Plot of accuracy as a function of RT bins (conditional accuracy) using same dataset as Fig. 1. Data pooled across all stimulus luminance and mice (n=8), sorted by stimulus size; RT bin size = 100 ms. Solid curves: Conditional accuracy functions (CAFs, best-fit rising asymptotic function; Methods) for targets of different sizes (black: 45⁰; blue: 35⁰; red: 25⁰); light shading: 95% CI of the fit (Methods). Histograms at bottom: RT distributions for targets of different sizes (y-axis on the right). The overall response accuracy for a particular stimulus condition is the dot product of the CAF and the RT distribution. **(B)** Box plots of the key parameters for different target sizes. Left panel: a_peak_; middle panel: slope parameter; right panel: t_peak_. **(C)** CAFs for targets of different luminance conditions (magenta: ‘low’ luminance - first three luminance levels from Fig. 1C; green: ‘high’ luminance - last four luminance levels; Methods); conventions as in A. **(D)** Box plots of the key parameters for different luminance conditions; conventions as in C. The box plots in all panels show the median (open circle), the 25th and 75th percentiles (the bottom and top edge of the box), and the most extreme data points not considered as outliers (whiskers); in some panels, the boxes are the same size as the symbol for the median. See also Fig. 2-1, 2-2.

**Figure 3.**
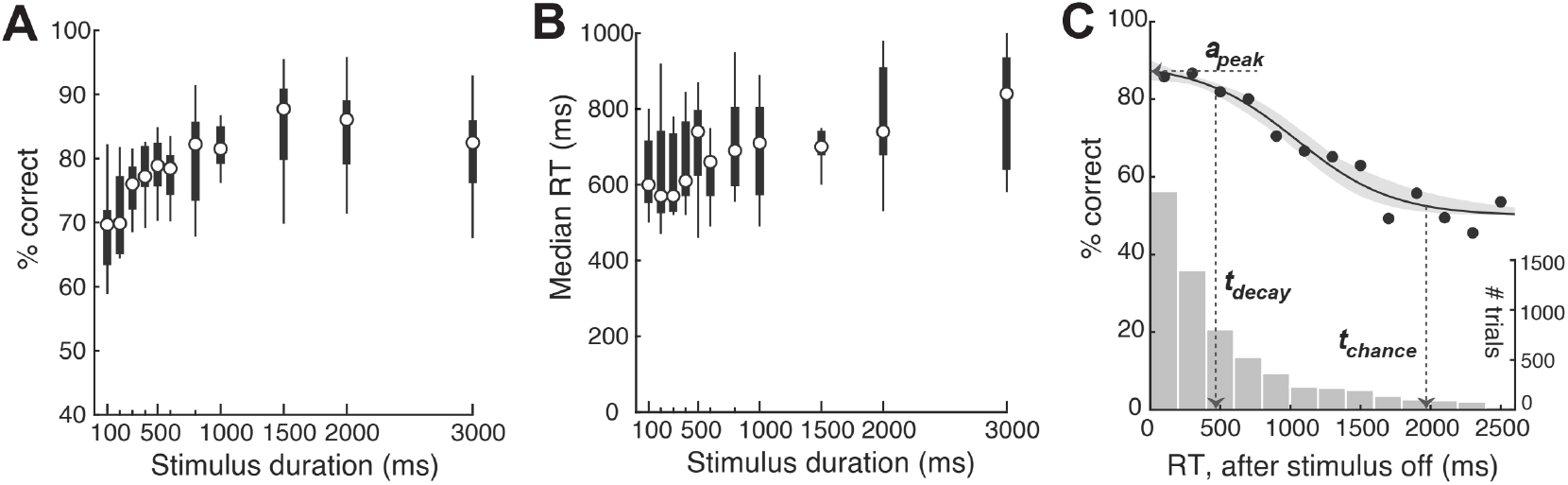
Stimulus duration and the memory-dependent stage of the conditional accuracy function. **(A)** Psychometric plot of discrimination accuracy against stimulus duration (n=9 mice; 1-way ANOVA; p<0.001. effect size η^2^=0.331). **(B)** Plot of median reaction time (RT) against stimulus duration (1-way ANOVA; p=0.056. effect size η^2^=0.177). **(C)** Plot of the conditional accuracy (solid data) as a function of RT bins relative to stimulus offset. Only trials in which the stimulus was longer than 332 ms were included (in order to ensure full sensory encoding - see text; Methods). Curve and shading: best-fit sigmoid function and 95% C.I. Bootstrapped estimates of each key metric: a_peak_, median [C.I.] =87.3 [84.8, 89.9] %; t_decay_ = 469 [279, 697] ms; and t_chance_ = 1969 [1708, 2520] ms. Histogram: RT distribution (y axis on the right). In this experiment, stimulus size and luminance were maintained fixed at 25°and 130 cd/m^2^respectively. See also Fig. 3-1.

### Flanker task

Upon trial initiation, either one stimulus (‘target’, 60 pix × 60 pix, luminance = 20.1 cd/m^2^, Michelson’s contrast=88%) was presented at the lower location, or two stimuli were presented simultaneously, with the target at the lower location and a second ‘flanker’ at the upper location (Fig.4A). Flankers were of the same size (60 pix × 60 pix) and spatial frequency (24 pixel/cycle) as the target, but with luminance ranging (over 8 levels) from less than that of the target to greater than that of the target [24]. The orientation of the flanker was either identical to that of the target (‘congruent trial’) or orthogonal to that of the target (‘incongruent trial’). The stimulus (stimuli) was (were) presented for a duration of 1s, and mice were required to report orientation of the target grating with the appropriate nose-touch (within 3s). All types of trials (no flanker, congruent, incongruent) and flanker contrasts were interleaved randomly within each daily session. Data from this experiment have been reported previously [24], and were re-analyzed here using different analyses.

### Subject inclusion/exclusion

A total of 37 mice were used in this study, with different subsets used in different tasks. For mice involved in more than one task, they were well-rested for 3-8 weeks with food and water *ad libitum* between experiments. Before the start of each experiment, all mice were given a few days of practice session to ensure that they remembered/re-learned the association between the orientation of single target and the appropriate nose-touch. Of the total of 37 mice trained across tasks, 28 mice passed the inclusion threshold of response accuracy >70% in the single stimulus discrimination task, and were included for the analyses reported in this paper.

### Trial inclusion/exclusion

Mice were observed to become less engaged in the task towards the end of a behavioral session, when they had received a sizeable proportion of their daily water intake. This was reflected in their behavioral metrics: they tended to wait longer to initiate the next trial, and their performance deteriorated. We identified and excluded such trials following a published procedure [24], in order to minimize confounds arising from loss of motivation towards the end of sessions. Briefly, we pooled data across all mice and all sessions, treating them as coming from one session of a single ‘mouse’. We then binned the data by trial number within the session, computed the discrimination accuracy in each bin (% correct), and plotted it as a function of trial number within session (Fig. 1-1B, 3-1A, 5-1A). Using a bootstrapping approach, we computed the 95% confidence interval for this value. We used the following exclusion criterion: Trials q and above were dropped if the q^th^ trial was the first trial at which *at least one* of the following two conditions was satisfied: (a) the performance was statistically indistinguishable from chance on the q^th^ trial and for the majority (3/5) of the next 5 trials (including the q^th^), (b) the number of observations in q^th^ trial was below 25% of the maximum possible number of observations for each trial (Σ mice*sessions), thereby signaling substantially reduced statistical power available to reliably compare performance to chance. The plots of performance as a function of trial number, and number of observations as a function of trial number for the different tasks in this study are shown in Figs. 1-1B, 3-1A, 5-1A, along with the identified cut-off trial numbers (q).

### Behavioral measurements

Response accuracy (% correct) was calculated as the number of correct trials divided by the total number of trials responded (correct plus incorrect). Reaction time (RT) was defined as the time between the start of stimulus presentation and time of response nose-touch, both detected by the touchscreen. In the experiment involving stimulus onset delays (Fig. 5A), RT was computed with respect to trial initiation (as opposed to from stimulus onset).

### Drift diffusion modeling of RT distributions

The RT measured here represents the duration from stimulus onset to completion of execution of the motor response. In order to specifically isolate the time spent in decision making (separately from the latency of activation of sensory neurons as well as duration of motor execution), we applied the drift-diffusion model to our RT data [40, 41]. This model hypothesizes that a subject (‘decision maker’) collects information from the sensory stimulus via sequential sampling, causing sensory evidence to accrue for or against a particular option (usually binary) while viewing the stimulus. A decision is to be made when the accumulating evidence reaches an internal threshold of the subject. This process of evidence accumulation, together with the processes of sensory encoding and motor execution, as well as threshold crossing, determine the RT observed on each trial.

We used a standard version of the model that consists of four independent variables [38, 42]: (1) the drift rate, (2) the boundary separation, (3) the starting point, and a (4) non-decisional constant (t_delay_), which accounts for the time spent in sensory encoding and motor execution. In the case of our tasks, there was no reason for the drift rate to be different between vertical versus horizontal gratings, and therefore, we merged both type of trials (trials with a horizontal target grating and trials with a vertical target grating). We treated ‘correct’ response and ‘incorrect’ response as the two binary options, and fit the diffusion model to the RT distributions of correct versus incorrect trials using the fast-dm-30 toolbox with the maximum likelihood option to gain estimates of those four parameters for each individual mouse (Fig. 2-2)[40].

### Conditional accuracy analysis

Conditional accuracy was calculated as the percentage of correct trials (accuracy) as a function of RT. For this analysis, trials from all mice were pooled together and treated as if they were from one single mouse for statistical power (Fig. 2 onwards; for completeness, conditional accuracy plots using non-pooled data, i.e., from individual mice, are included in Extended Figures). Pooled trials were then sorted by their RT, and then binned by RT such that there were: (1) sufficient number of trials in each bin; and (2) sufficient number of total bins, to ensure the robustness of curve fitting and therefore the estimates of quantitative metrics (see below). Typical bin sizes used were 50ms, 100 ms or 200 ms bins, depending on the experiments and stage of analysis (sensory encoding or STM-dependent). The effect of bin size on the estimates of quantitative metrics is explored in the Extended Figures; results show that the estimates are comparable across tested bin sizes.

### Conditional accuracy function (CAF)

To quantitatively describe the relationship between the conditional accuracy and RT, we fitted the plot of accuracy against binned RT with parametric functions (the CAF; see below) using a nonlinear least square method For RT bins aligned to stimulus onset (Fig. 2, 4C, 5B), we fit the conditional accuracy data using an increasing asymptotic function:

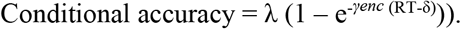

**Figure 4.**
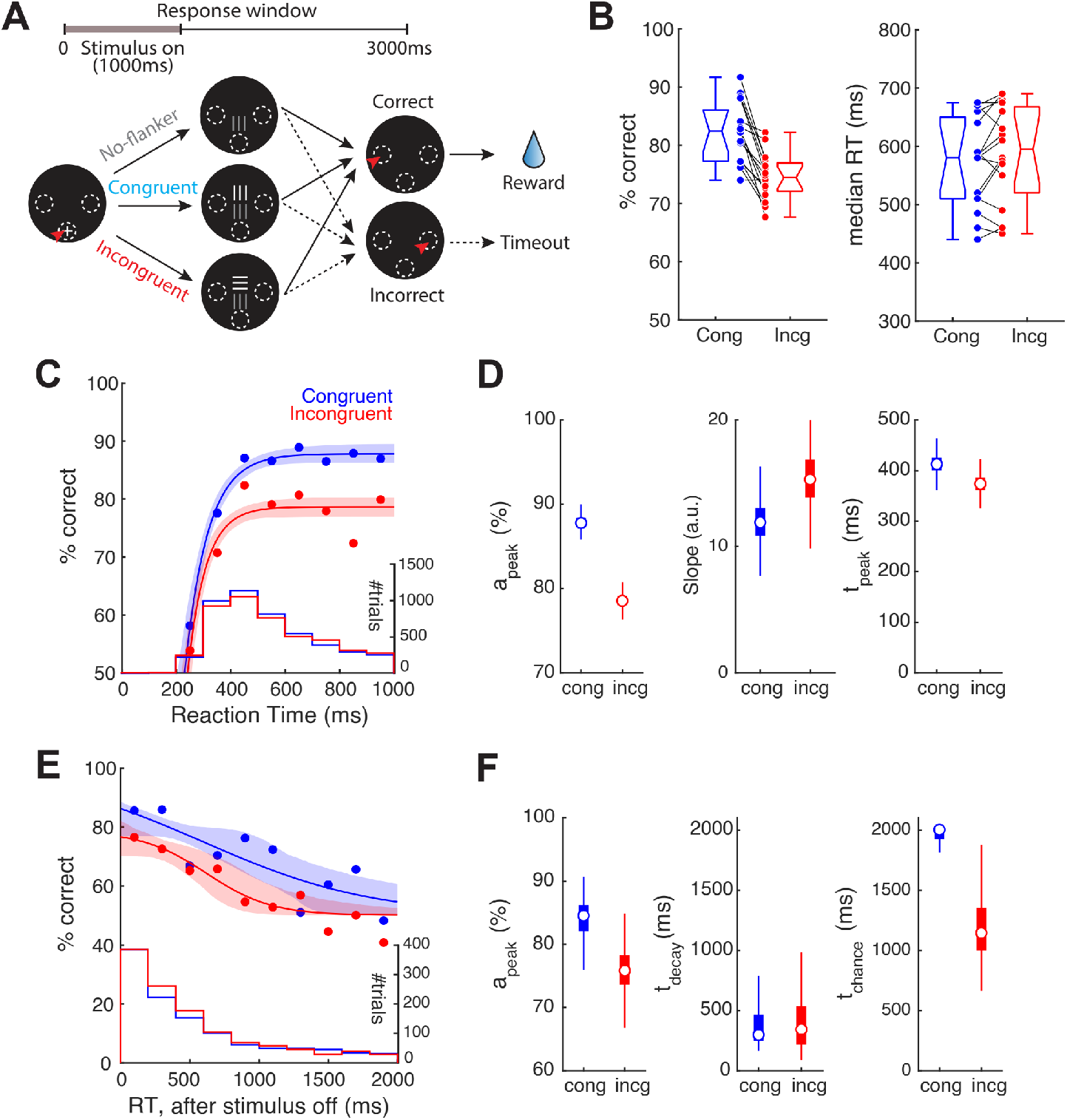
Incongruent flanker modulates the sensory encoding stage of the conditional accuracy function (CAF). **(A)** Schematic of the flanker task; target grating is always presented at the lower location; a second ‘flanker’ grating (orthogonal orientation – incongruent flanker, or same orientation – congruent flanker) is presented simultaneously, and always at the upper location; contrast of flanker is systematically varied (adapted from [24]). All other conventions as in Figure 1. The stimuli were presented for 1s and the response window was 3s. **(B)**<u>Left panel</u>: Comparison of performance between trials with incongruent vs. congruent flanker. p<0.001, paired-sample *t* test. effect size Hedges’ g=1.61. <u>Right panel</u>: Comparison of median RT between trials with incongruent vs. congruent flanker. p=0.137, paired-sample *t* test. effect size Hedges’ g=−0.176. Data re-analyzed from You et al [24]; each line represents data from one mouse (n=17 mice). Data in B-F include only trials with high flanker luminance (≥20.1 cd/m^2^; see text). **(C)** CAFs of the sensory encoding stage; Blue: trials with congruent flanker; red: trials with incongruent flanker; histograms; RT distributions. **(D)** Key parameters of CAFs (sensory encoding stage) for trials with congruent vs. incongruent flanker; a_peak_ (left), slope parameter (middle), and t_peak_ (right). Box plots show the distribution of bootstrapped estimates (Methods). Effect sizes (congruent – incongruent): a_peak_: Hedges’ g=11.0; slope parameter: Hedges’ g=−1.73; t_peak_: Hedges’ g=2.08. Note, the sizes of the boxes in the left and right panels are similar to the sizes of the circular symbols depicting the medians. **(E)** CAFs of the STM-dependent stage; data aligned to stimulus offset. Blue: trials with congruent flanker; red: trials with incongruent flanker. **(F)** Plots of key parameters of CAFs (STM-dependent stage) for trials with congruent vs. incongruent flanker; a_peak_ (left), t_chance_ (middle) and t_decay_ (right). Conventions and statistical methods as in D. a_peak_: Hedges’ g=2.54; t_chance_: Hedges’ g=2.98; t_decay_: Hedges’ g=0.175.

**Figure 5.**
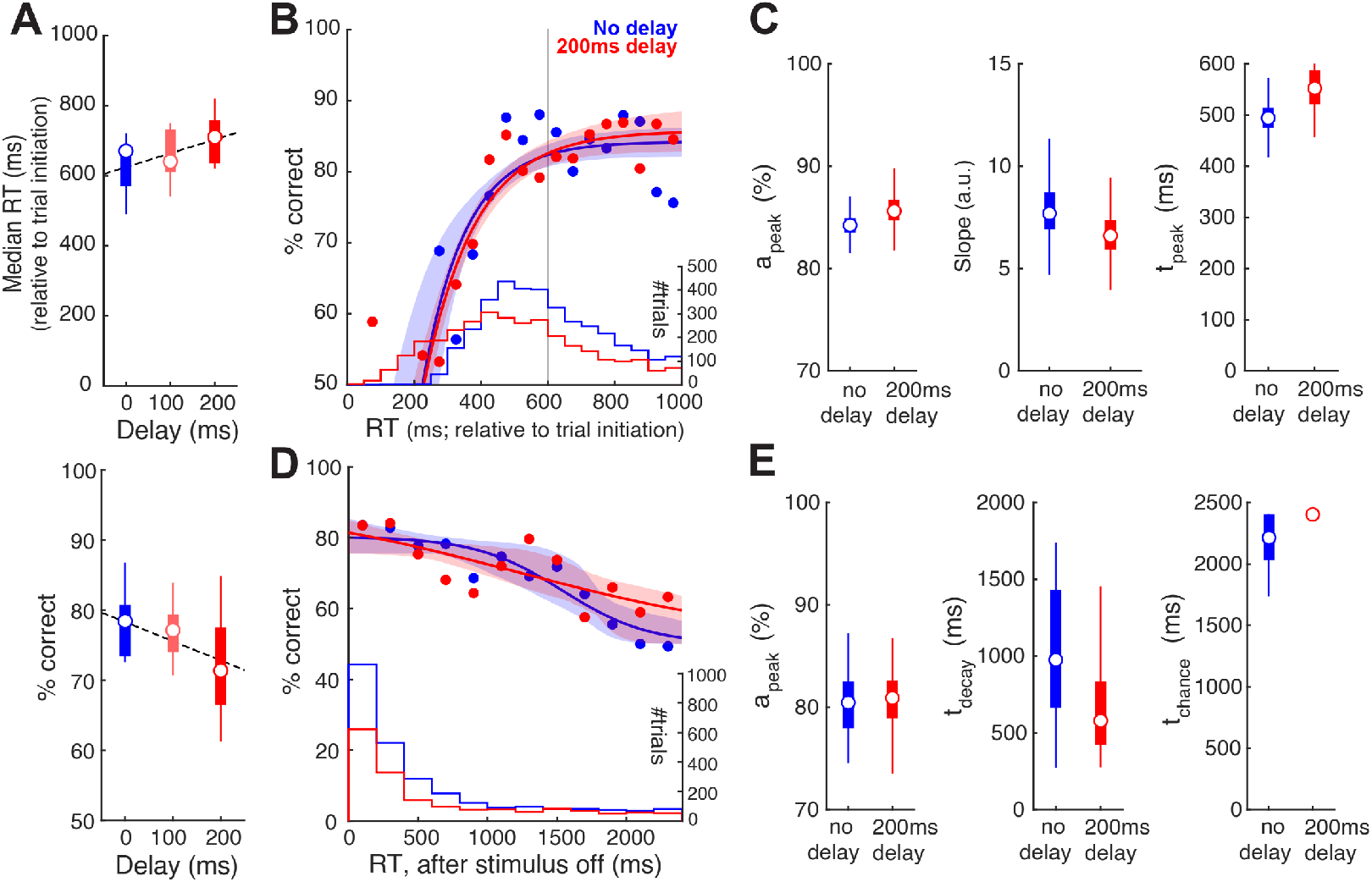
Stimulus onset delay modulates RT distribution but not the conditional accuracy function. **(A)**<u>*Upper:*</u> Plot of median RT, measured relative to train initiation, against stimulus onset delay (n=9 mice; p=0.094, 1-way ANOVA; effect size η^2^=0.179; Pearson’s correlation=0.422, p=0.028). Dashed line: Linear regression on RTs. <u>*Lower:*</u> Plot of response accuracy against stimulus onset delay (p=0.182, 1-way ANOVA; effect size η^2^=0.132; Pearson’s correlation=−0.358, p=0.067). **(B)** Conditional accuracy functions of the sensory encoding stage; Blue: trials with no delay; red: trials with 200ms delay; shaded bands: bootstrap confidence intervals (95%); confidence intervals overlap for the two datasets. Histograms: RT distributions. Grey vertical line: stimulus offset. **(C)** Key parameters of the CAF (sensory encoding stage) for trials with no delay vs. trials with 200ms delay. Box plots show the distribution of the bootstrapped estimates. **(D)** Conditional accuracy functions of the STM-dependent stage. Conventions as in B. (E) Key parameters of the CAF (STM-dependent stage) for trials with no delay vs. trials with 200ms delay. Conventions as in C. **See also Fig. 5-1.**

Three key metrics were defined for this sensory encoding phase, for use in subsequent comparisons across conditions: (1) peak conditional accuracy (*a_pea_*_k_), the maximal level of accuracy that the CAF reaches within the range of RT; (2) the slope parameter (*γ_enc_*); and (3) the first instant at which the conditional accuracy reaches its maximal value (t_peak_) - defined as the time point at which the ascending CAF reaches 95% of *a_pea_*_k_. Note that t_peak_ is influenced by the peak conditional accuracy (*a_peak_*, the slope parameter, *γ_enc_*, and the temporal offset at chance performance, δ For RT bins aligned to stimulus offset (Fig. 3C, 4E, 5D), we fit the decaying conditional accuracy data using a sigmoidal function:

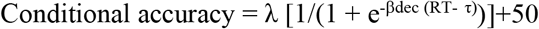

Three key metrics were defined for this STM-dependent stage for use in subsequent comparisons across conditions: (1) peak conditional accuracy (*a_pea_*_k_), the maximal level of accuracy within the range of RT; (2) the first instant (t_decay_) at which conditional accuracy is lower than the maximum - defined as the time point at which the descending CAF crosses 95% of *a_pea_*_k_; and (3) the first instant (t_chance_) at which conditional accuracy drops to chance levels - defined as the timepoint at which the descending CAF crosses 52.5%. In (rare) cases when the CAF never went below 52.5%, t_chance_ was set to be the upper bound of the window of analysis (i.e., 3000ms – stimulus duration = the window for which the mice can respond following stimulus offset). Note that t_decay_ and t_chance_ are influenced by both the slope parameter, β_dec_, and τ.

Confidence intervals of the CAF fits, as well as for the parameters, were estimated by standard bootstrapping procedures involving resampling the raw data randomly with replacement (1000 x), to get repeated estimates of the CAF and corresponding metrics. In all relevant figures, the box plots of the estimated values of each metric show the median (the central mark), the 25th and 75th percentiles (the bottom and top edge of the box), and the most extreme data points not considered as outliers (whiskers).

In the experiment in which the stimulus onset delay was manipulated (Fig. 5), we adopted the following two adjustments to our procedure for the analysis of the conditional accuracy function. First, since the stimulus was short (600 ms), in order to ensure robust estimates of CAF metrics for the sensory encoding stage, we included data beyond stimulus offset as well for the fitting of the CAF through 400 ms following offset. (We chose to include data upto 400 ms after offset, specifically, because we had learned from Figure 3 that conditional accuracy remains at its plateau for nearly 500 ms following stimulus offset.) Second, we also excluded trials with RT < 200ms for the fitting of the CAF (Fig. 5B), because these represent trials on which responses were initiated prematurely (200 ms represents our estimate of the duration of sensory latency plus motor execution; see text surrounding Figure 2).

### Statistical tests

All analyses and statistical tests were performed in MATLAB. For single-stimulus experiments in which only one stimulus parameter was systemically varied, one-way ANOVA was applied to examine the effect of the manipulating the single factor (duration and delay, Fig. 3AB, 5A, 1-1CD). For experiments that involved changing both stimulus size and contrast (Fig. 1CDE, 2-2), two-way ANOVA was applied to examine the effect of each factor, as well as their interaction. For the flanker task, the paired-sample t-test was used to examine if the group performance was different between trial types (Fig. 4B).

For the metrics associated with CAF, comparisons were made by measuring the effect size (Hedges’ g) of the difference between two distributions (Fig. 2BD, 4DF and 5CE). All effect size measurements, including those with ANOVA (η^2^), were calculated following the methods (and source code) of Hentschke and Stȕttgen (2011)[43]. Hedges’ g estimates the distance between the two distributions in units of their pooled standard deviation, with larger numbers indicating stronger effects. η^2^ varies from 0 to 1, with larger values indicating greater ratio of variance explained in the dependent variable by a predictor while controlling the other variables.

## RESULTS

In this study, freely behaving mice were trained to perform 2-AFC orientation discrimination in a touchscreen-based setup [24, 29](Methods). Briefly, mice were placed in a plexiglass tube within a soundproof operant chamber equipped with a touch-sensitive screen at one wall and a reward well at the opposite wall (Fig. 1A). A plexiglass sheet with three holes was placed in front of the touchscre–n - the holes corresponded to the locations at which mice were allowed to interact with the screen by a nose-touch (Fig. 1A). All trials began with a nose-touch on a bright zeroing-cross presented within the lower central hole (Fig. 1B). Immediately following nose-touch, an oriented grating (target; bright stimulus on a dark background) was presented at the center of the screen. Mice were rewarded if they responded to the orientation of the target with an appropriate nose-touch: vertical (horizontal) grating → touch within upper left (upper right) hole. Behavioral data were collected from daily sessions that lasted 30 minutes for each mouse.

### Stimulus size and luminance modulate mouse discrimination performance

We first examined the effect of target size and target contrast on the decision performance of mice in the orientation discrimination task. Here, the target grating was presented for up to 3 seconds after trial initiation (Fig.1B; Methods), and its size and luminance were systematically varied; the spatial frequency was fixed at 0.1 cycles/degree (24 pixels/cycle) [16, 17] (Methods). Mice were allowed to respond at any time during stimulus presentation, and the stimulus was terminated automatically upon response.

We found that both the target contrast and size significantly modulated discrimination accuracy (Fig. 1C, 2-way ANOVA, main effect of luminance, p<0.001, effect size η^2^=0.292; main effect of size, p<0.001, η^2^=0.192; interaction, p=0.498, η^2^=0.037). These results revealed that mice discriminated target orientation better than chance even at the lowest luminance (2.00 cd/m^2^) and size (25°) tested (Fig. 1C; the red box at the left lower corner, p=0.015, *t*-test against mean accuracy=50%, effect size g1=1.129). Additionally, at this smallest target size (25⁰), mice could discriminate with >80% accuracy for most of the tested luminance values (≥4.37 cd/m^2^; Fig. 1CD, red data).

The effect of these parameters on median reaction times (RTs) was less pronounced. Target size, but not contrast, modulated reaction times (RTs) (Fig.1E, two-way ANOVA; main effect of size, p=0.004, effect size η^2^=0.071; main effect of luminance, p=0.998, η^2^=0.003; interaction, p=1, η^2^=0.010). Together, these results revealed a systematic effect of target size and luminance on discrimination accuracy.

### Effect of stimulus size and contrast on dynamics of visual decision-making: the sensory encoding stage

To investigate the dynamics of visual perceptual decision-making, we adapted approaches from human studies and examined the dependence of response accuracy on RT, i.e., the so-called ‘conditional accuracy’ function (CAF) [9–11]. For these analyses, we pooled trials from all mice (n=8) in order to gain better statistical power for the estimates of parameters of the CAF (Methods; plots of the data for individual mice showed similar overall shapes of the CAF; Fig.2-1A).

Specifically, we investigated the dynamics of visual perceptual decision-making as a function of stimulus size, and separately, as a function of stimulus luminance. First, to examine the effect of stimulus size on decision dynamics, we pooled trials from all mice across luminance values (7 luminance values) for each stimulus size, sorted them based on RT, and plotted conditional accuracy for each RT bin (100ms; Fig. 2A; Methods). We found that for responses with RT less than ~500 ms, conditional accuracy improved for longer RT, consistent with the classic ‘speed-accuracy tradeoff’ [34]. For responses with RT greater than 500 ms and up to 3s, the allowed duration for responses, conditional accuracy plateaued, and was independent of RT. Next, to examine the effect of stimulus luminance on decision dynamics, we pooled trials from all mice across size values into two groups based on stimulus luminance: (1) trials with target luminance ≤ 4.37 cd/m^2^ (‘low luminance’), and (2) trials with target luminance > 4.37 cd/m^2^ (‘high luminance’; Methods). Here, as well, we found a similar initial stage of increasing conditional accuracy upto RT of ~ 500 ms, followed by a plateauing of conditional accuracy.

Drawing upon arguments from human behavioral studies, we reasoned that the initial transient stage of the conditional accuracy function reflects the process of sensory encoding: during it, slower responses allow more sensory evidence to be acquired, thereby improving conditional accuracy up to a peak value reflecting the completion of sensory encoding [30–33].

To quantify these dynamics, we fit the conditional accuracy data with an asymptotic function (Fig. 2AC, solid curves) [9–11], and estimated three key metrics, in each case: (1) the peak conditional accuracy (*a_pea_*_k_), (2) the slope parameter (γ*enc*), and (3) the timepoint at which conditional accuracy reached its peak (*t_peak_;* Methods).

We found that the peak conditional accuracy was significantly modulated by stimulus size (Fig.2B-left; *a_peak_*: size 25°, median [C.I.] = 81.3 [79.1, 83.7] %; size 35⁰ = 88.0 [86.5, 89.4] %; size 45⁰ = 92.4 [90.7, 94.1] %; effect size Hedge’s g= −6.71 (25°-35°), −5.39 (35°-45°), −10.6 (25°-45°)), but not the slope of the function (slope parameter, γ*enc*, Fig. 2B-middle, size 25° = 6.52 [5.10, 9.07] a.u.; size 35° = 8.81 [7.09, 10.6] a.u.; size 45° = 7.92 [6.15, 10.1] a.u. Hedges’ g= −2.24 (25°-35°), 0.863 (35°-45°), −1.34 (25°-45°)), or the time to reach peak accuracy (t_peak_, Fig. 2B-right, size 25° = 493 [375, 597] ms; size 35° = 459 [420, 505] ms, size 45° = 466 [420, 522] ms; Hedges’ g= 0.728 (25°-35°), −0.274 (35°-45°), 0.558 (25°-45°)).

Next, we found that the peak conditional accuracy was higher in high-luminance trials (Fig. 2D-left, low-luminance, median [C.I.] = 84.7 [82.7, 86.3] %; high-luminance = 89.5 [88.2, 90.7] %, effect size Hedges’ g= −6.13). The slope was also higher in high-luminance trials (slope parameter, γ*enc*, Fig. 2D-middle, low-luminance = 6.37 [5.21, 7.78] a.u.; high-luminance = 10.32 [8.49, 12.6] a.u., Hedges’ g= −4.51) suggesting a faster rate of sensory encoding in high-luminance trials. Consistent with this, the time to reach peak accuracy was shorter in high-luminance trials (Fig.2D-right; t_peak_: low-luminance = 531 [478, 599] ms; high-contrast = 412 [378, 448] ms, Hedges’ g= 4.86).

The RT measured here represents the duration from the start of the sensory input to the completion of execution of the motor response. In order to obtain an estimate of the duration, specifically, of decision-making, we employed the standard the drift diffusion modeling (DDM) approach [38, 42] (Methods). Briefly, the DDM analyzes the full RT distribution and yields a quantitative estimate of four parameters (Methods), one of which is t_delay_, a parameter which accounts for the combination of: (a) the time taken for the sensory (visual) periphery to transduce and relay information to visual brain areas (i.e., neural response latency), as well as (b) the time taken for executing the motor response (i.e., motor execution delay). In our tasks, the latter corresponds to the time for the mouse to move its head (and body) to achieve the appropriate nose-touch.

Using this approach, we found that stimulus size as well as luminance had no discernable effect on t_delay_ (Fig. 2-2. 2-way ANOVA, size: p=0.308, luminance: p=0.523; interaction: p=0.931), and the average value of t_delay_ was 212 ms. Consequently, we estimated the duration of just the sensory encoding stage (temporal integration window) as t_peak_ – t_delay_ = t_peak_ – 212 ms. Across conditions, this took values of 200 ms (412 ms −212 ms; high luminance), 254 ms (466-212 ms; size of 45 deg), 247 ms (459-212 ms; size of 35 deg), 281 ms (493-212 ms; size of 25 deg), and 319 ms (531 ms −212 ms, low luminance).

Thus, conditional accuracy analysis allowed us to quantify the sensory encoding stage in mouse visual perceptual dynamics. We estimated its duration to be brief, varying between 200 ms and 320 ms across the tested conditions.

Following the completion of sensory encoding, a fully constructed representation of the sensory stimulus is available, as a result of which, additional sampling of the stimulus brings no extra benefits to the performance. Our finding that RTs longer than t_peak_ produce no further increase in conditional accuracy, is consistent with the view (Fig. 2AC).

### Stimulus duration and the dynamics of visual decision-making: the memory-dependent stage

The next stage in the time course of perceptual decisions has been identified in human studies as the so-called ‘short-term memory’ (STM)-dependent stage, during which an internal representation of the sensory stimulus is available transiently in memory for guiding behavior [44]. Studies have demonstrated the STM to be labile such that once the stimulus is terminated, sensory information maintained in STM decays and is lost (over seconds) [45–49].

In our experiments so far, the target stimulus was present on the screen for the full duration of the response window (3s). Here, in order to investigate and quantify the STM-dependent stage of mouse perceptual decisions, we performed an experiment in which we shortened the stimulus duration systematically from 3s to 100 ms. This allowed us to examine the time course of decision behavior following stimulus offset, and, as well, to examine the shortest stimulus that mice are able to discriminate effectively.

We first examined overall mouse behavioral performance at different stimulus durations. We found that response accuracy was significantly modulated (Fig.3A, one-way ANOVA, p<0.001, effect size η^2^=0.331), with accuracy decreasing for shorter stimulus durations (Pearson’s ρ=0.712, p=0.014). There was also a trend of decreasing median RT for shorter stimulus durations (Fig.3B, one-way ANOVA, p=0.056, effect size η^2^=0.177; Pearson’s ρ=0.861, p=0.001). Additionally, these results revealed, that the shortest stimulus duration needed for mice to be able to discriminate above chance was less than 100 ms - the smallest duration tested (Fig. 3B).

Next, to examine the decision dynamics following stimulus offset, we aligned trials to stimulus offset, and computed the conditional accuracy. Considering that incomplete sensory encoding may be a confounding factor to the STM decay, we only included those trials on which the stimulus was presented for longer than the duration of the sensory encoding stage, estimated in Figure 2 to be 320 ms.

We observed the classic decay in conditional accuracy with longer RTs (Fig. 3C). To quantify the time course of the decay, we fit the conditional accuracy data with a sigmoidal function (Methods), and estimated three key metrics (Fig. 3C; Methods). The first, peak performance, a_peak_, was 87.3% (median, C.I.= [84.8, 89.9] %), comparable to the asymptotic level of Figure 2, thereby supporting that sensory encoding is, indeed, complete on these trials. The second, the time point at which the conditional accuracy dropped below the peak value, t_decay_, was 469 ms (median, C.I.= [279, 697] ms) after stimulus offset. The third, the first timepoint at which the discrimination accuracy dropped to a level indistinguishable from the chance, t_chance,_ was 1969 ms (median, C.I.= [1708, 2520] ms) after stimulus offset (Methods).

Thus, our conditional accuracy analysis allowed us to investigate quantitatively the second, STM-dependent stage in mouse visual perceptual dynamics. We estimated the duration over which above-chance decision accuracy is supported in mice after stimulus offset as ~1700ms (i.e., t_chance_ minus the t_delay_).

### The presence of flanker stimulus modulates perceptual dynamics

We next investigated the impact of sensory context on visual decision dynamics. It is well-established that the sensory context in which the perceptual target is presented modulates animals’ behavior [50–52]. For instance, in the classic flanker task in humans, the co-occurrence of a flanker stimulus with conflicting information can interfere with perceptual performance [53, 54]. Recently, similar results were demonstrated in mice using a touchscreen version of the flanker task [24]. In this task (Fig. 4A), a target grating (always presented at the lower location) was accompanied by a flanker grating at the upper location with either orthogonal orientation (‘incongruent’ flanker) or same orientation (‘congruent’ flanker). Compared to the presence of a congruent flanker, the ‘incongruent’ flanker significantly impaired discrimination accuracy (Fig. 4B-left; p<0.001, paired-sample *t* test. effect size Hedges’ g=1.61; re-plotted based on data from [24]; Methods). Here, we analyzed that dataset with the conditional accuracy analysis to investigate whether an incongruent flanker affected the sensory encoding stage or the STM-dependent stage of perceptual dynamics.

To investigate the effect of the flanker on perceptual dynamics, we pooled trials from all mice into two groups based on their flanker congruency, and sorted the trials based on their RT. Since previous study [24] has demonstrated that the flanker affects performance significantly only when its luminance is higher than (or equal to) that of the target, here we included only high-luminance trials (trials with flanker luminance ≥20.1 cd/m^2^). To examine the sensory encoding stage quantitatively, we followed the approach used in Figure 2 and selected the trials on which mice responded before the stimulus ended (RT < 1000ms), and aligned them to stimulus onset. Separately, to examine the STM-dependent stage, we followed the approach used in Figure 3 and selected the trials on which responses were made after the stimulus ended, and aligned them to stimulus offset.

The sensory encoding stage was significantly modulated by flanker congruency (Fig. 4CD). We found that, the peak conditional accuracy for incongruent trials was significantly lower than that of congruent trials (Fig. 4D-left; *a_peak_*: congruent, median [C.I.] = 87.8 [86.3, 89.6] %, incongruent = 78.5 [76.9, 80.2] %; effect size (congruent-incongruent) Hedges’ g=11.0), indicating that the presence of a high-luminance incongruent flanker interfered with the sensory encoding of the target stimulus. While the slope parameter for incongruent trials remains comparable to that of the congruent trials (Fig. 4D middle; congruent = 11.9 [9.10, 15.5] a.u., incongruent = 15.3 [11.6, 20.6] a.u.; Hedges’ g=−1.73), the time to reach peak accuracy was, however, shorter for incongruent trials (Fig. 4D-right; *t_peak_:* congruent = 413 [378, 458] ms, incongruent = 374 [340, 410] ms; Hedges’ g=2.08), consistent with the lower *a_peak_* (Fig. 4D-left).

The STM-dependent stage also appeared to be modulated by flanker congruency (Fig. 4EF). Following stimulus offset, the time at which conditional accuracy dropped to chance was much earlier in incongruent trials than in congruent trials (Fig. 4F-right; *t_chance_*: congruent, median [C.I.] = 2000 [1363, 2000] ms; incongruent = 1145 [816, 1985] ms; Hedges’ g=2.98). However, this was likely due largely to the lower peak conditional accuracy for incongruent trials (Fig.4F-left; *a_peak_*: congruent= 84.5 [76.9, 88.6] %; incongruent = 75.9 [70.1, 82.1] %; Hedges’ g=2.54), as opposed to changes in t_decay_ (Fig. 4F-middle; congruent= 299 [197, 1086] ms; incongruent= 343 [126, 802] ms; Hedges’ g=0.175), or to the rate of decay (slope parameter; data not shown, congruent= −1.82 [−10, −1.0] a.u., incongruent= −4.82 [−10.0, −1.60] a.u.; Hedges’ g=0.99).

In sum, we found that the presence of an incongruent flanker interferes the sensory encoding stage but not the STM-dependent stage of mouse visual decision dynamics.

### Stimulus onset delay modulates RT distribution but not the conditional accuracy function

The components of behavioral performance that we have investigated thus far, namely, overall decision accuracy, RT distribution and conditional accuracy function are related formally in the following way: the overall decision accuracy is the dot product of the conditional accuracy function and RT distribution.

Our manipulations, thus far, produced changes in the conditional accuracy function predominantly. Here, we wondered whether task parameters could, instead, alter RT distribution, and possibly do so without affecting the conditional accuracy function. To test this, we added a delay between trial initiation and target onset (called stimulus onset delay) in the single stimulus discrimination task. We reasoned that the extent to which mice are unable to adaptively withhold responding could impact the RT distribution.

We found that adding a stimulus onset delay did alter the RT distribution of mice (Fig. 5A-upper panel). The median RTs, measured relative to trial initiation, showed an increasing trend with delay (one-way ANOVA, p=0.094; effect size η^2^=0.179; Pearson’s correlation=0.422, p=0.028). This indicated that mice were able to sense the delayed onset of stimulus and thereby withhold their responses. However, mice were unable to withhold responding for the full duration required. By performing a linear regression (Fig. 5A-upper panel; dashed line), we found that mice were able to withhold their responses for only 39 ms for every 100ms of delay. Separately, this increase in RT for longer delays was accompanied by a trend towards lower decision accuracy (Fig. 5A-lower panel, one-way ANOVA, p=0.182; effect size η^2^=0.132; Pearson’s correlation=−0.358, p=0.067).

By contrast, conditional accuracy analysis revealed no effect of stimulus onset delay either on the sensory encoding stage (Fig. 5BC, *a_peak_*: no-delay, median [C.I.] = 84.2 [82.2, 86.4]%, 200ms-delay = 85.6 [82.9, 89.2]%, effect size (no-delay - 200ms-delay) Hedges’ g=−1.12; slope parameter: no-delay = 7.69 [5.82, 10.7] a.u., 200ms-delay = 6.61 [4.63, 8.74] a.u., Hedges’ g=0.264; t_peak_: no-delay = 494 [436, 557] ms, 200ms-delay = 552 [476, 680] ms, Hedges’ g=−1.49), or on the STM-dependent stage (Fig. 5DE, *a_peak_*: no-delay, median [C.I.] = 80.5 [75.6, 85.5]%, 200ms-delay = 80.9 [75.6, 84.9]%, Hedges’ g=−0.147; t_decay_: no-delay = 976 [332, 1642] ms, 200ms-delay = 580 [319, 1585] ms, Hedges’ g=0.877; t_chance_: no-delay = 2214 [1865, 2400] ms, 200ms-delay = 2400 [1935, 2400] ms, Hedges’ g=−1.22).

Taken together, our results from varying the stimulus onset delay show that changes in RT distribution (and overall decision accuracy) are not necessarily accompanied by changes in the conditional accuracy function. The observed trend of decreased accuracy was accounted for by the fact that with a delay, there were more responses initiated before the sensory encoding was complete, or even before the stimulus was presented (i.e., ‘impulsive’ responses) (Fig.5B, histograms). To quantify such impulsivity, we propose an ‘impulsivity index’ (ImpI): ImpI = 1 – average (duration for which mice withhold responses /duration of the delay). Higher positive values of this index indicate greater impulsivity, with ImpI=1 indicating a complete inability to withhold responding in the face of stimulus delays (‘maximally’ impulsive). In the case of our mice, ImpI is ~0.6.

## DISCUSSION

In this study, we quantify two distinct stages in the temporal dynamics of visual perceptual decisions in mice. First, a sensory encoding stage that is subject to the speed-accuracy tradeoff, and then, a short-term memory dependent stage in which decision performance decays once the stimulus disappears. We also demonstrate that the conditional accuracy function and the RT distribution can be affected independently by experimental manipulations. Whereas stimulus size, luminance and presence of a foil modulate the conditional accuracy function with minimal changes to the RT distribution, stimulus onset asynchrony modulates the RT distribution without changes to the conditional accuracy function. Additionally, our results yield numerical estimates of fundamental psychophysical constants of visual perceptual decision-making in mice. Taken together, this study establishes a quantitative platform for future work dissecting neural circuit underpinnings of the dynamics of visually guided decision-making in mice.

### Estimates of time constants of the dynamics of visual perceptual decision-making in mice

Our results yielded numerical estimates of the duration of sensory encoding (i.e., the window of temporal integration) as 200-320 ms across stimulus size and luminance (Fig. 2). This estimate is similar to that in humans: the internal representation of a visual stimulus is thought to be constructed within the first 200-300 ms of stimulus presentation [30–33]

On the other hand, we also obtained an estimate of the duration of STM as 1700 ms. This constituted the period starting from stimulus offset to the last instant at which responses that are better than chance were *initiated* (Fig. 3D; t_chance_ - t_delay_ = ~1700 ms). This duration does not necessarily represent just the maintenance of visual stimulus information in STM, it could also represent maintenance of information about the motor response associated with the stimulus (and likely, a combination of the two). Notably, our estimate of the duration of viability of the labile internal representation in mice falls in the same range as has been reported from human studies [34, 35, 55].

We have interpreted the decay in performance following stimulus offset as being due to loss of information in STM. A potential confounding factor to this interpretation is differences in the internal state of the animal – in selective attention, or more generally, task engagement. It is possible, for instance, that all the trials with longer RTs represent those in which mice did not pay attention to the stimulus (or more generally, were disengaged from the task), thereby also being associated with lower accuracy. Indeed, in our flanker task, we find that disruption of attention interferes with sensory encoding and causes the conditional accuracy to be lower following stimulus offset (Fig. 4).

However, unlike in the flanker task, in these trials, attention was not varied systematically, suggesting that a loss of attention (or more generally, changes in internal state) are unlikely to account systematically for the late decay of conditional accuracy. (Indeed, if they did, that would predict that with a steady level of attentiveness or engagement, response accuracy would never decay following stimulus offset, in direct contravention to published literature.) Nonetheless, because it is difficult to quantify the extent to which changes in internal state may have played a role in our task, we propose that our estimate of the duration of STM of 1700 ms serves as a *lower bound* for the duration of STM.

This estimate of 1700 ms also represents a lower bound for working memory (WM). Whereas STM refers to the retention of information even when it is not accessible from the environment, WM is thought of as ‘STM+,’ referring additionally to the ability to manipulate this information and protect it from interference [56, 57]. WM can be lengthened with training. For instance, in tasks that require animals hold information over an enforced delay period before responding, it has been reported that mice can learn to perform well with delay periods up to 5 sec [58]. Here, by allowing the natural evolution of the dynamics of decision-making to occur without an imposed delay period, we were able to estimate the ‘intrinsic’ (lower bound for the) duration of STM.

### Estimates of the operating range of stimulus features for visual perceptual decision-making in mice

This study also yielded estimates for the range of values of various stimulus features within which mice are able to discriminate the visual target. The smallest stimulus and lowest luminance at which mice were able to discriminate orientation above chance were 25° and 2.00 cd/m^2^, with mice performing at > 80% accuracy for most luminance values at that smallest size. The shortest stimulus that mice are able to discriminate above chance was ≤ 100ms (Fig. 3A). Further, based on the x-intercept of the CAF in sensory encoding stage (median [C.I.] = 236 [215, 253] ms, pooling all trials of various sizes and luminance from Fig. 2), we were able to refine this estimate to be ≤ 53ms (conservatively, after subtracting t_delay_ = ~200 ms). This is consistent to the estimation (40-80 ms) from a previous study based on visual cortical activity [59]. In a subgroup of animals (n=3), we tested if mice are able to discriminate orientation of the target stimulus (25°, 0.1cpd, 16.2 cd/m^2^) when it was 50 ms long. Two out of the three mice showed a response accuracy higher than chance (accuracy = 57.9%, 210 correct out of 363 trials, p=0.002, binomial test; and 55.6%, 143/257, p=0.040, respectively), consistent with this refined estimate. These findings that mice are able to discriminate visual stimuli in demanding sensory contexts suggest that the visual perceptual abilities of mice may be underrated.

The best discrimination performance reported in mice (accuracies > 90%) have typically been obtained using large, often full-field, grating stimuli [18, 60]. In our single target discrimination task, the best performance ranged lower, between 75-90% (Fig. 1C), consistent with our use of ‘small’ stimuli (relative to those typically used in mouse vision studies [15, 16, 18, 20, 61]) and the lower visual acuity of mice. Indeed, in our pilot study, the performance plateaued at ~93% for a stimulus size ≥ 45° (Fig. 1-1CD). These results suggest that full-field stimuli may be effectively replaced by 45° stimuli to achieve best performance levels.

The best discrimination performance exhibited a dip at the highest luminance (Fig. 1C). This is potentially well accounted for by signal saturation: because the visual system adapts to the relevant range of stimulus luminance for best encoding [62], the interleaved presentation of stimuli with different luminance can render the maximum-luminance stimulus unfavorable because of signal saturation [18]. Consistent with this idea, when the maximum-luminance stimulus (25°, 0.1cpd, 130 cd/m^2^) was presented *alone* in block design (Fig. 1-1C, the green box at the left most, group median [C.I.] = 85.7 [77.6, 92.1] %), response accuracy was nominally higher than when it was presented interleaved with stimuli of varying luminance (Fig. 1C, the red box at the right most, group median [C.I.] = 79.7 [61.9, 91.9] %). These results indicate that a good upper bound for stimulus luminance in mouse experiments may be ~34 cd/m^2^.

### Stimulus and task parameters modulate perceptual performance through a variety of mechanisms

Increase in stimulus size and luminance both improve the discrimination performance of mice (Fig. 1). However, analysis of conditional accuracy revealed that increasing the stimulus size and luminance both increased the peak conditional accuracy (a_peak_), but only increasing the stimulus luminance increased the slope of the CAF and resulted in a shorter t_peak_ (Fig. 2). We propose that these differences in the CAF actually reflect differential mechanisms underlying the processing of stimulus size versus *contrast*, as opposed to stimulus size versus *luminance*. On the one hand, in our experiments, varying luminance (by varying the intensity of the bright phase of the grating), also varied stimulus contrast (relative to the dark background). On the other, increasing stimulus size increased the total luminance while maintaining the contrast fixed. Consequently, the observed differences in the CAF following manipulations of stimulus size and luminance are best explained by differential mechanisms for stimulus size versus contrast processing.

Separately, manipulating attention (by presenting a flanker) and the stimulus onset asynchrony both caused a reduction in response accuracy (Fig.4B, 5A-lower panel). However, again, the analysis of conditional accuracy suggests that the mechanisms underlying the two are different: the capture of attention by the flanker interferes with the target’s sensory encoding, whereas adding a pre-stimulus onset delay results in change of the RT distribution without affecting the CAF.

Taken together, our results demonstrate that although manipulating stimulus parameters or experimental conditions may induce similar changes in perceptual performance (overall accuracy), their underlying mechanisms could be very different. The conditional accuracy analysis serves as an informative tool to investigate these mechanisms in detail and to understand the dynamics of perceptual decision making.

### Qualitative differences between perceptual decision-making tasks as well as between task-difficulties

Our results, in conjunction with published studies, suggest that qualitatively different visual perceptual decision-making tasks may produce sensory encoding stages with substantially different time scales. Across the various tasks and stimulus conditions that we studied here in mice, the sensory encoding stage ended rapidly around 300 ms, exhibiting an asymptotic relationship between conditional accuracy and RT. However, in a recent study in which rats discriminated the direction of motion of a patch of randomly moving dots, the sensory encoding stage continued through 1.5 s (the longest RT bin reported), exhibiting a linear relationship between conditional accuracy and RT, and pointing to an even longer encoding stage [63]. We propose that this large difference in the duration of sensory encoding may be due to fundamentally different natures of the tasks: tasks involving noise and stimulus dynamics (as with the random dot motion patch) may necessitate longer time windows for sensory integration, compared to tasks with non-noisy, static stimuli (as with all of our tasks).

Our results also highlight that ‘task difficulty’ may be altered in qualitatively different ways, producing distinct outcomes on behavior. In the literature, task difficulty is often increased by making target stimuli noisier, or more ambiguous, or by introducing distracters (which we did also). Such manipulations often cause subjects (animals) to respond slower, allowing them time to either gather more information to produce better performance. We found similar results here as well. Additionally, we shortened the stimulus duration, which can plausibly be considered to also increase task difficulty. However, when we did so, we found the opposite result – mice responded faster as the target stimulus became shorter (Fig. 3B). This potentially counter-intuitive effect (faster RTs for a ‘more difficult’ task) is explained well by the conditional accuracy analysis (Fig. 3C). Whereas shortening the stimulus duration makes the task more difficult, responding more slowly to shorter stimuli does not grant a perceptual benefit to the animals: once the stimulus has disappeared, withholding responses for longer would only increase the risk of losing information owing to memory decay. In other words, short stimuli impose a ‘time pressure’ on animals to make decisions quickly. Thus, task difficulty may be altered in qualitatively different ways, with distinct behavioral effects.

### Optimal sensory sampling during visual perceptual decision-making in mice

An intriguing observation in our study is that across tasks, the peak of RT distribution always seemed to occur around t_peak_ (Fig. 2AC, 4C). Since the RT distribution can vary independently of the conditional accuracy function (as demonstrated in Fig. 5), there is no *priori* reason that the peak of RT distribution (or median RT) and the t_peak_ must change together. We propose that responding with RTs close to t_peak_ is, in fact, an optimal behavioral strategy for the mice. As indicated by the conditional accuracy function, mouse response accuracy increased as RT increased until it reached a plateau at t_peak_. Responding earlier than t_peak,_ therefore, would sacrifice accuracy, while responding later than t_peak_ would needlessly delay response (reducing the reward rate). Consequently, responding with the peak of RT distribution being equal to t_peak_ would be optimal.

## EXTENDED DATA

Extended data (Fig. 1-1, 2-1, 2-2, 3-1, and 5-1) and legends are included.

## Extended Data

**Figure 1-1. Extended data for Figure 1.**
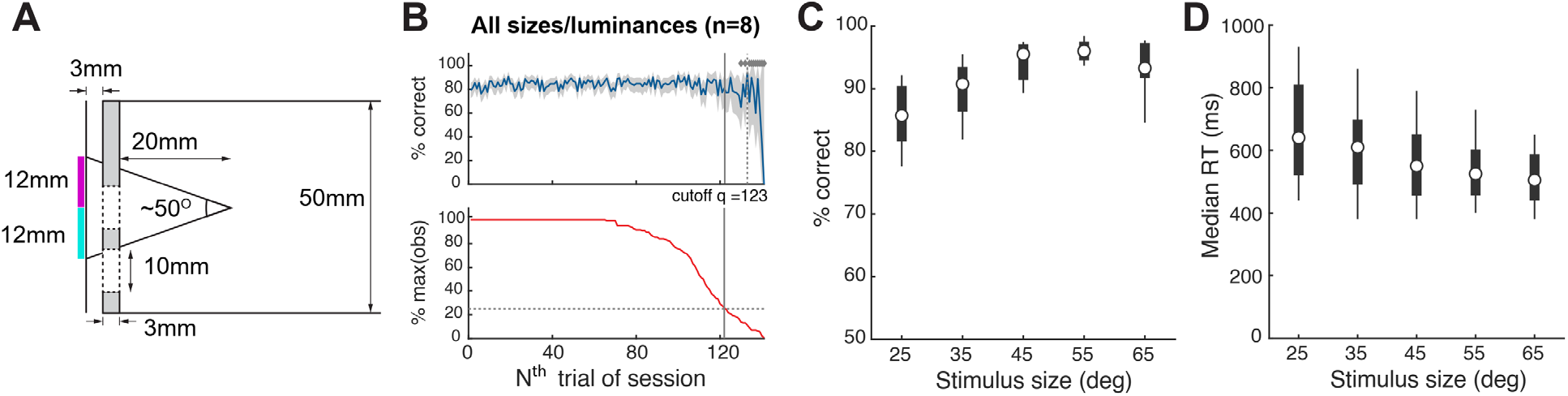
**(A)** Lateral view of the schematic experimental setup showing the relative position of the touchscreen (leftmost vertical line), the plexiglass mask (grey-filled vertical bar), and the tube within which mice move (50 mm diameter); the plexiglass mask is positioned 3 mm in front of the touchscreen. Dashed lines indicate the central response hole (lower dashed lines), and left/right response holes (upper dashed lines; 10 mm diameter). For single-stimulus discrimination, the center of the stimulus is aligned with the center of left/right response holes in elevation, and with the central hole in azimuth (see Fig.1A). For experiments involving two stimulus locations (i.e., flanker task), the upper (magenta) and lower (cyan) locations of the stimulus are indicated as colored bars (see also Fig. 4A). The 60 pixels × 60 pixels (12mm × 12mm) stimulus subtends a visual angle of 25⁰ when viewed from 20 mm front of the plexiglass mask. **(B)** Identification of trials towards the end of the 30 min behavioral sessions that corresponded to animals being poorly engaged in the task (Methods and [24]). <u>Top panel</u>: Time course of overall response accuracy across mice as a function of trial number within sessions. Accuracy obtained from trials pooled across all mice and sessions, and computed as a function of trial number within session (blue; Methods). Grey shading: bootstrapped estimates of the 95% confidence interval of the accuracy (gray; Methods). Diamonds on top: trials whose accuracy not significantly different from chance. Dashed vertical line: first trial at which the accuracy was not different from chance (50%), and stayed indistinguishable from chance for 3/5 of the next 5 trials (Methods). Data show increased variability and worse performance towards the end of sessions. <u>Bottom panel</u>: Number of actual observations across mice for each trial number, as a percentage of the maximal number of possible observations (Σ mice*sessions), plotted as a function of trial number within session (red). Solid vertical line: first trial at which the number of observations drops below 25%. Data show drop in the number of observations available to reliably assess performance towards the end of sessions. Based on these data, all trials above 122 of each behavioral session of this experiment were dropped from analysis (Methods). Results in Fig. 1 are based on data from trials 1-122 from each behavioral session. (**C)** Response accuracy as a function of stimulus size (n=9 mice; p=0.001, 1-way ANOVA). In these experiments, stimulus size was manipulated independently (without manipulation of luminance; unlike in Figure 1). All stimuli were at the highest luminance (130 cd/m^2^, or Michelson-contrast of 98%). **(D**) Median RT as a function of stimulus size (n=9 mice; p=0.205, 1-way ANOVA).

**Figure 2-1. Extended data for Figure 2:**
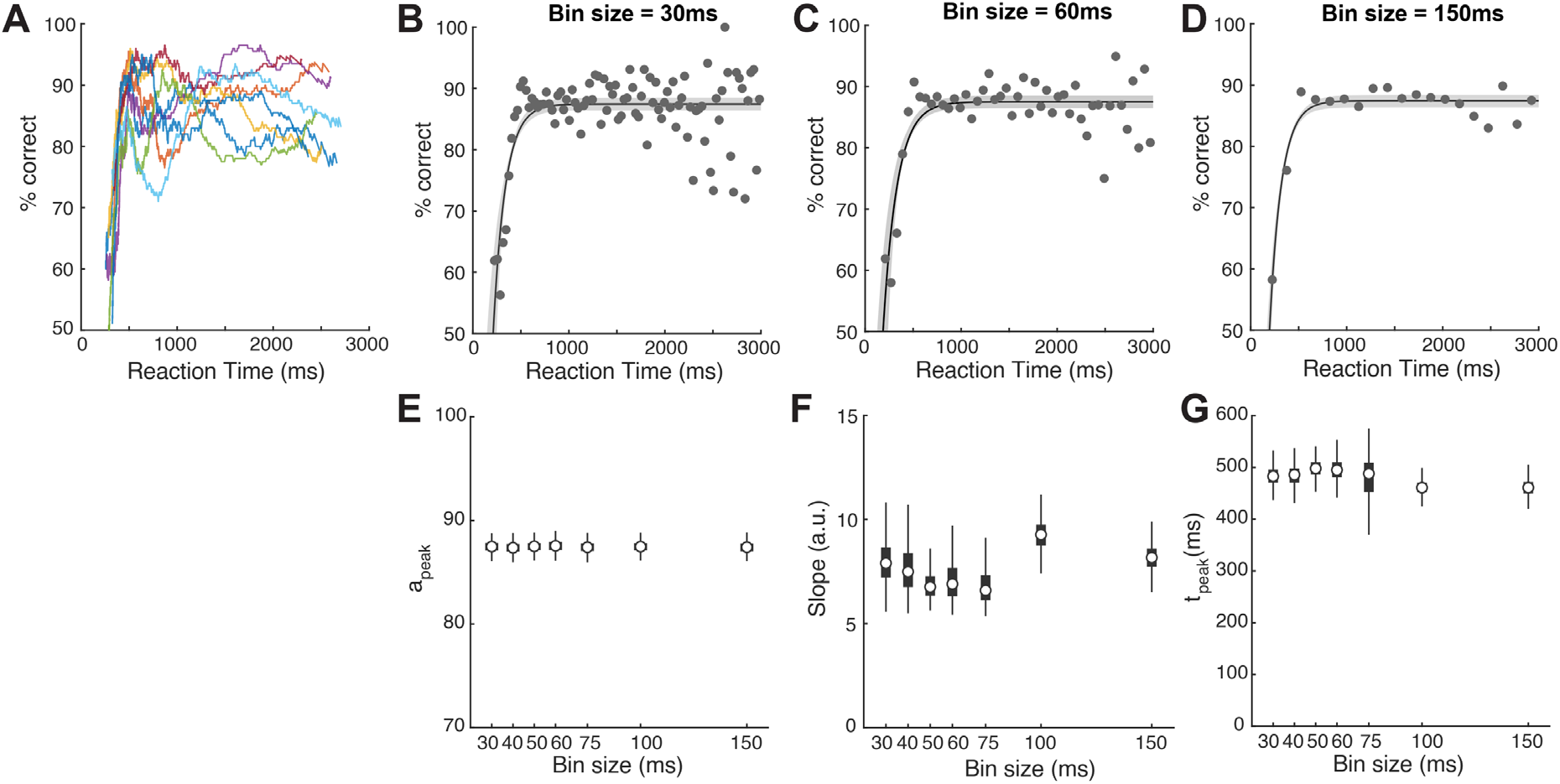
CAFs of individual mice and effect of bin size. **(A)** The general pattern of conditional accuracy curves across mice. Each color represents one single mouse. Each curve was generated by pooling all trials (of various stimulus size and luminance) from one mouse, sort the trials by RT, and then do a moving average (window size = 200 trials) to plot the mean accuracy (y) at mean RT (x) of the time window. **(B-D)** Fitting of the conditional accuracy function (CAF) in various bin sizes. (B) Bin size = 30ms; (C) Bin size = 60ms; (D) Bin size = 150ms; **(E-G)** Estimates of the quantitative metrics of the CAF in various bin sizes. (E) peak conditional accuracy (a_peak_); (F) slope parameter; and (G) time to reach peak conditional accuracy (t_peak_).

**Figure 2-2. Extended data for Figure 2:**
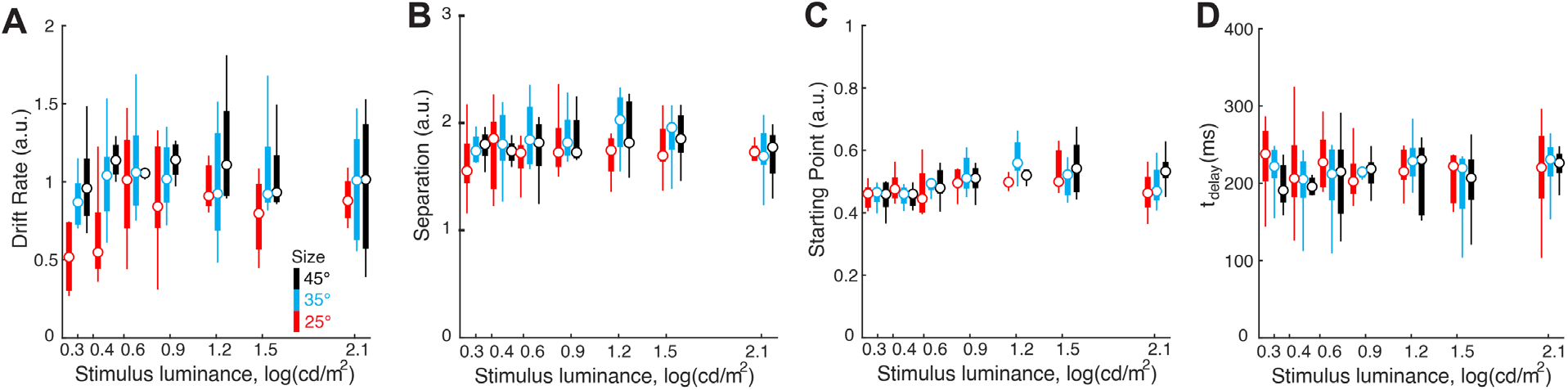
Estimates of all four parameters of the drift diffusion model. **(A)** Drift rate; 2-way ANOVA, p=0.028 (luminance), p<0.001 (size), p=0.767 (interaction). **(B)** Boundary separation; 2-way ANOVA, p=0.171 (luminance), p=0.026 (size), p=0.953 (interaction). **(C)** Starting point; 2-way ANOVA, p<0.001 (luminance), p=0.325 (size), p=0.098 (interaction). **(D)** t_delay_; 2-way ANOVA, p=0.523 (luminance), p=0.308 (size), p=0.931 (interaction).

**Figure 3-1. Extended data for Figure 3:**
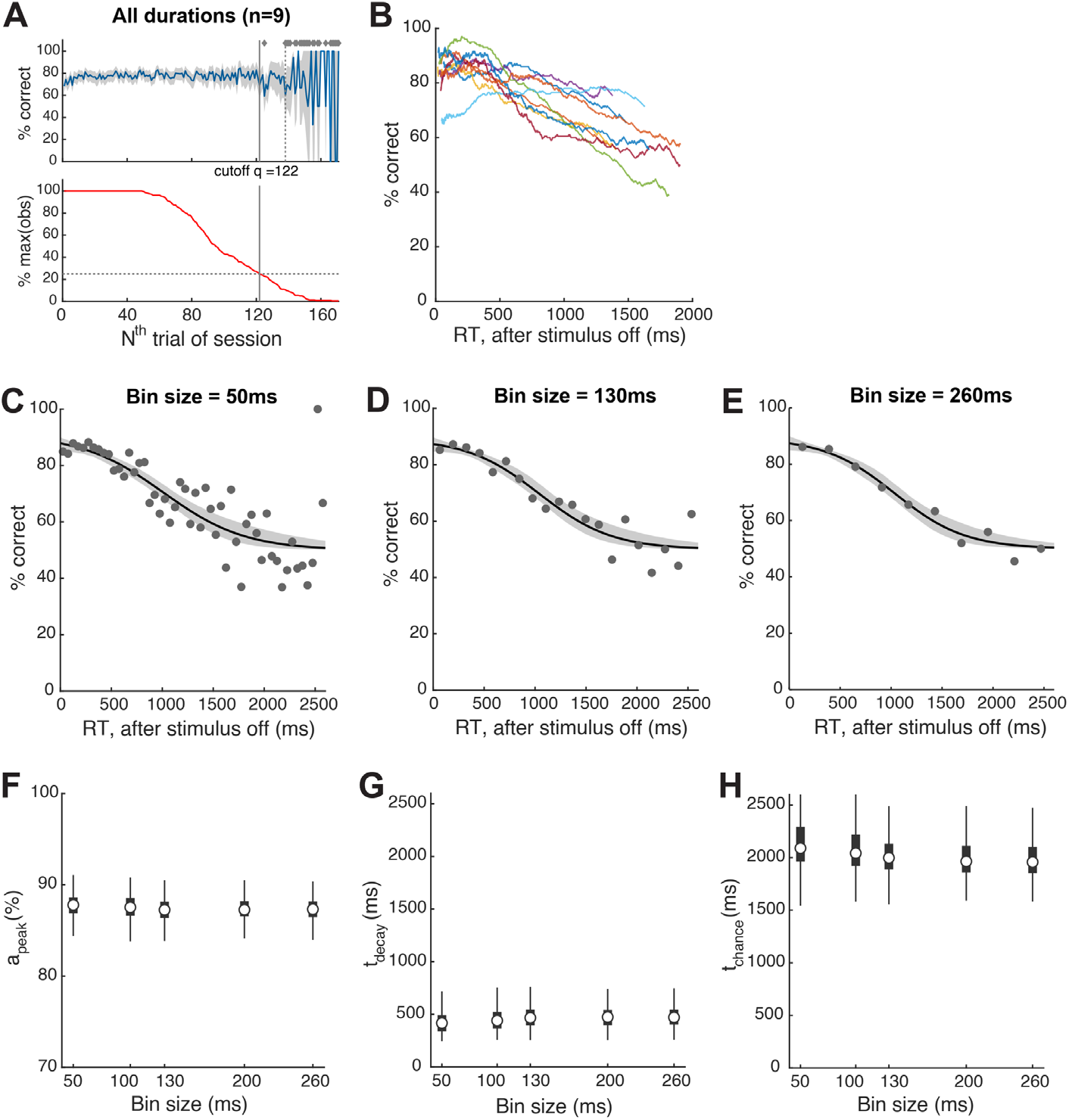
CAFs of individual mice and effect of bin size. **(A)** Identification of trials towards the end of the 30 min behavioral sessions that corresponded to animals being poorly engaged in the task (Methods); conventions identical to those in Fig.1-1B. **(B)** The general pattern of conditional accuracy curves across mice. Each color represents one single mouse. Each curve was generated by pooling all trials (of various stimulus size and luminance) from one mouse, sort the trials by RT, and then do a moving average (window size = 200 trials) to plot the mean accuracy (y) at mean RT (x) of the time window. **(C-E)** Fitting of the conditional accuracy function (CAF) in various bin sizes. (C) Bin size = 50ms; (D) Bin size = 130ms; (E) Bin size = 260ms; **(F-H)** Estimates of the quantitative metrics of the CAF in various bin sizes. (F) peak conditional accuracy (a_peak_); (G) the time at which conditional accuracy started to decay (t_decay_); and (G) the time at which conditional accuracy fell to the chance level (t_chance_).

**Figure 5-1.**
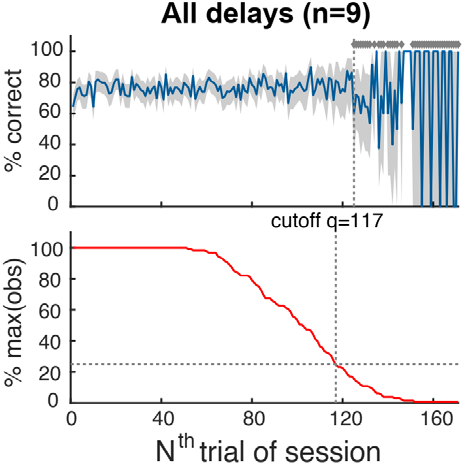
Extended data for stimulus onset delay experiment. Identification of trials towards the end of the 30 min behavioral sessions that corresponded to animals being poorly engaged in the task (Methods). All conventions are as in Fig.1-1B. Based on these data, all trials above 116 of each behavioral session of this experiment were dropped from analysis. Results in Fig.5 are based on data from trials 1-116 from each behavioral session.

